# Why the Sleeping Brain Clears

**DOI:** 10.64898/2026.04.16.718904

**Authors:** Christian Kerskens

**Affiliations:** Trinity College Institute of Neuroscience

## Abstract

The mechanical origin of cerebrospinal fluid (CSF) and interstitial fluid (ISF) transport remains unresolved. High-frequency arterial pulsations (∼ 1 Hz) have long been proposed as a driver of CSF flow, yet multiple biomechanical analyses suggest that their ability to support deep bulk interstitial transport is severely limited by the poroelastic resistance of neural tissue. At the same time, slow-wave sleep is associated with large, synchronous CSF oscillations and enhanced clearance-related dynamics near ∼ 0.05 Hz. What selects this low-frequency regime remains unclear.

Here we propose a theoretical framework in which this frequency selection is not incidental, but mechanically necessary. When neural populations update their state, local thermodynamic demand induces microvascular dilation. Under intracranial volume constraints, this blood-volume expansion must, to leading order, be compensated by displacement of other intracranial volume components, including CSF. We model the poroelastic response of the interstitial matrix and obtain an effective low-pass filter for this displacement, with a nominal cut-off frequency in the slow-wave range (*r*_*c*_ ≈ 0.05 Hz).

This mechanical filter implies two distinct forcing regimes. During wakefulness, rapid commitment and sensorimotor resetting are hypothesized to generate spectrally sharp, high-frequency transients in vascular volume. Because this spectral content lies largely above the poroelastic passband, waking dynamics are predicted to be inefficient at driving deep bulk transport. Slow-wave sleep, by contrast, reduces rapid commitment-like transitions and permits smoother, more globally synchronized vascular-volume oscillations that fall within the passband and support larger-scale CSF motion.

The framework yields several falsifiable predictions, including load-dependent modulation of sleep-associated CSF pulsation amplitudes, a BOLD-first / CSF-second temporal ordering during slow-wave events, and a mechanical discrepancy between deep interstitial transport and the rapid dispersion of superficial exogenous tracers. More generally, the theory advances a strong claim: the sleeping brain is mechanically privileged for large-scale CSF dynamics not because sleep introduces a new driver, but because sleep permits forcing in a frequency range that brain tissue can actually transmit.

## Introduction

The brain must solve two physical problems at once: it must process information and it must clear metabolic waste. The second problem is especially acute because the brain lacks a dedicated lymphatic pump, yet must remove large molecular byproducts, including amyloid-beta and tau, through cerebrospinal fluid (CSF) and interstitial fluid (ISF) transport [1, 2]. Although substantial progress has been made in characterizing perivascular and glymphatic pathways, the dominant mechanical drivers of large-scale fluid transport, and the extent to which clearance relies on directed bulk flow versus passive diffusion [3, 4], remain fiercely debated.

A central difficulty is the mismatch between candidate driving forces and the observed state dependence of large-scale clearance-related dynamics. Arterial pulsations (∼ 1 Hz) have long been proposed as a driver of CSF flow [5], but rigorous theoretical and biomechanical analyses suggest that such high-frequency forcing is strongly attenuated by the poroelastic and geometrically tortuous structure of neural tissue before it can support deep bulk interstitial transport [6, 7]. Respiratory forcing [8] and intrinsic vasomotion [9, 10] provide additional candidate drivers, but they do not by themselves explain why the largest macroscopic CSF oscillations and strongest clearance-related signatures appear during slow-wave sleep, when global fluctuations emerge near ∼ 0.05 Hz [11, 12]. The interpretation of these mechanisms is further complicated by recent reports that exogenous tracers can disperse rapidly during wakefulness [13], a finding that appears difficult to reconcile with the strong sleep dependence reported for macroscopic endogenous waste clearance [14].

The central physical mechanism of this framework can be stated without initial reference to the geometric formalism. When neural populations update their state, local thermodynamic demand increases and the microvasculature dilates [15]. Under the Monro–Kellie doctrine [16, 17], intracranial volume is approximately fixed over short timescales. Because the skull is rigid, this blood-volume increase must displace a corresponding volume of CSF. If many brain regions dilate synchronously and slowly enough, the cumulative displacement becomes a brain-wide pump. The poroelastic resistance of neural tissue acts as a mechanical low-pass filter that selects which frequencies of this pumping can generate deep bulk flow. The information-geometric framework developed below formalizes this mechanism as a physiological manifestation of a mathematical triad connecting geometry, entropy, and substrate constraint [18]. This framework identifies a further distinction: the metabolic cost profile differs qualitatively between the sharp, commitment-driven dynamics of wakefulness and the smoother exploratory dynamics of sleep, which shifts the spectral content of the mechanical forcing.

The information-geometric framework developed below formalizes this mechanism as a physiological manifestation of a mathematical triad connecting geometry, entropy, and substrate constraint [18]. In this framework, the transition from wakefulness to sleep is interpreted as a geometric regime change: the high-frequency forcing associated with wakefulness is modeled as a stiffening of the effective metric, which reduces the mechanical compatibility of the substrate with deep bulk transport.

We formalize this proposal in three steps. First, we model cognitive state change as a gradient flow on a state space subject to finite perfusion capacity. This yields an effective Bures–Wasserstein metric back-reaction that we interpret physiologically as local cerebral blood-volume expansion. Second, we embed that volume change in the mechanics of a poroelastic tissue matrix and derive an effective low-pass filter for bulk fluid motion. Third, we use this filter to distinguish two regimes: a wakeful regime dominated by repeated boundary-approach forcing that is mechanically inefficient for clearance, and a slow synchronized sleep regime that can support macroscopic CSF transport. The full causal chain proposed here—from cognitive demand to vascular expansion, CSF displacement, and poroelastic filtering—is summarized schematically in Fig. 1.

**Figure 1:**
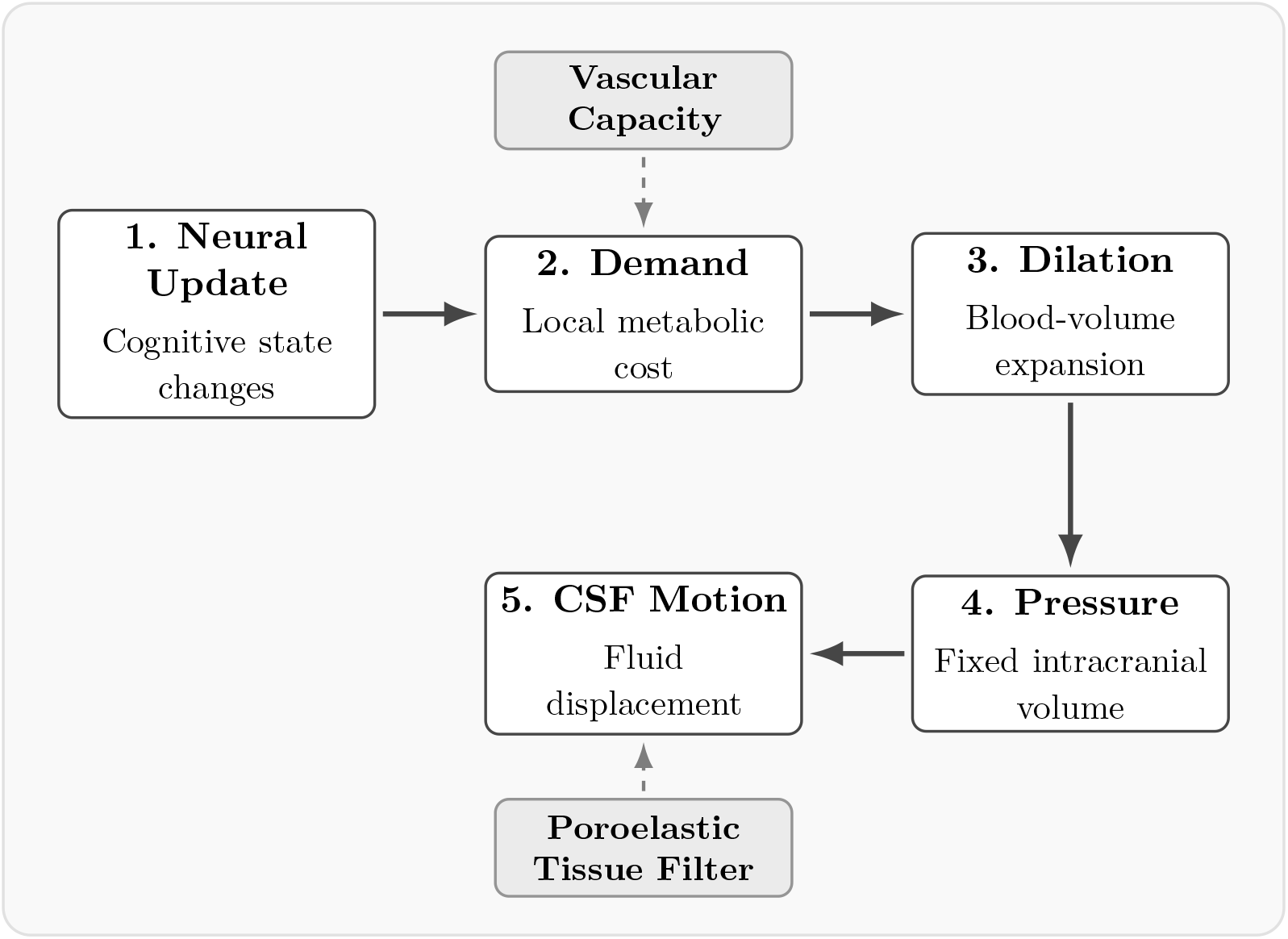
The causal chain of the thermodynamic CSF pump. The framework posits a five-step physical mechanism: (1) cognitive state updates drive (2) local thermodynamic demand, which is gated by finite vascular capacity to produce (3) vascular dilation. Under the (4) Monro–Kellie fixed-volume constraint, this expansion mechanically drives (5) oscillatory CSF motion. The efficiency of the resulting pump is selected by the poroelastic low-pass filter of the tissue matrix, favoring slow, synchronized oscillations.

This framework also offers a natural interpretation of the temporal ordering reported in sleep neuroimaging, in which a hemodynamic BOLD increase precedes the large CSF inflow pulse [11]. In the present model, the BOLD event reflects the vascular side of a coordinated metric back-reaction, whereas the delayed CSF wave is the mechanically filtered hydrodynamic consequence of that preceding blood-volume change. The slow CSF pulse is therefore not the primary event, but the delayed fluid response of a poroelastic intracranial system driven by synchronized vascular expansion.

On this view, the low frequency of sleep-associated CSF oscillations is not incidental. Rather, slow-wave sleep is mechanically privileged because it permits vascular forcing in a frequency range that brain tissue can actually transmit. The theory therefore offers a candidate mechanistic explanation for why clearance-related CSF dynamics are amplified during slow-wave sleep, while also generating quantitative predictions linking cognitive load, vascular capacity, hemodynamic timing, and CSF pulsation amplitude.

## Results

The mechanical argument in Sections 1–2 is self-contained: it establishes the poroelastic filter and its frequency-selection logic independently of any specific source model. Section 3 then introduces a geometric framework that specifies the source term and explains why wakefulness and sleep generate different forcing profiles. A field-theoretic formulation of the complete source-to-transport chain, in which the conformal response acts as an active-strain source and Monro–Kellie compensation emerges as a constrained boundary-flux law, is developed in **Supplementary Note S3**.

### 1. From cognitive demand to volume displacement

We begin by relating neural-state change to a local thermodynamic demand. In the framework used here, the effective neural state evolves on an information-geometric manifold under the gradient flow of a free-energy functional [19]. The instantaneous local cost of updating that state is taken to scale with the squared norm of the gradient-flow velocity,

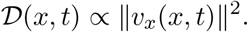

This proportionality is a constitutive modeling assumption, but it is a natural one. In the Benamou–Brenier formulation of optimal transport [20], the Bures–Wasserstein metric is defined through kinetic energy. In an overdamped biological medium, the power needed to drive state change against internal dissipation is correspondingly expected to scale with the square of the effective state-space velocity. In the present paper, 𝒟 (*x, t*) plays the role of the local thermodynamic load generated by inference.

The substrate, however, has finite capacity. Cerebral autoregulation can increase perfusion locally only up to a maximal effective capacity *Q*_max_(*x, t*). In the geometric framework developed in companion work, this capacity constraint modifies the effective Bures–Wasserstein metric through a self-consistent conformal factor. In the volumetric neurovascular case considered here, we take

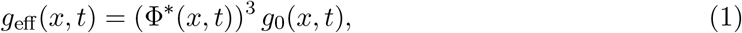

With

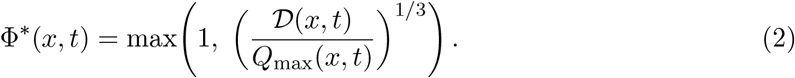

The cubic exponent reflects an isotropic volumetric interpretation of the local vascular response: the conformal stretching is read here as a space-filling local expansion of the perfused volume (which serves as a conservative lower bound when accounting for the fractal topology of the microvascular bed; see **Supplementary Note S1**). In that sense, the factor (Φ^*^)^3^ serves as a proxy for local cerebral blood-volume (CBV) expansion,

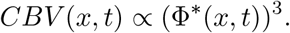

Under the Monro–Kellie doctrine, intracranial volume is approximately fixed over short timescales. A local increase in blood volume must, to leading order, be compensated by displacement of other intracranial volume components, including CSF and venous blood. In the simplified treatment adopted here, we focus on the CSF-displacement component. This gives the local mechanical relation

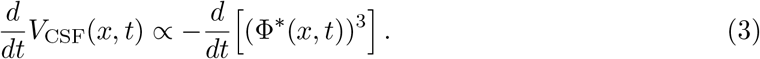

This is the core mechanical bridge of the paper. Thermodynamically constrained cognitive demand induces local vascular volume change, and that volume change induces CSF displacement. The Bures–Wasserstein metric modulation does not merely describe cognition abstractly; under the present physiological interpretation it provides a source term for fluid motion.

### 2. Poroelastic tissue response selects the slow-wave band

The preceding section identifies a candidate source of local fluid displacement. The next question is whether such displacement can drive bulk transport through brain tissue. The answer depends on the poroelastic response of the neural interstitium.

As detailed in the **Methods**, we model the tissue as a poroelastic medium in the sense of Biot theory [21, 22, 23]. The physical properties of this interstitial matrix, including its high tortuosity and strictly limited hydraulic permeability, are well established in the experimental and biomechanical literature [24]. Linearizing the response around a driven oscillatory regime yields an effective single-pole low-pass filter for macroscopic clearance:

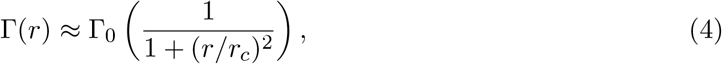

where *r* is the forcing frequency, Γ_0_ is the low-frequency clearance gain, and *r*_*c*_ is the poroelastic cut-off frequency.

Using conservative literature values for cortical hydraulic permeability and tissue stiffness— consistent with detailed 3D reconstructions of neuropil interstitial geometry [25]—we derive a nominal cut-off frequency *r*_*c*_ ≈ 0.053 Hz. This value lies directly in the slow-wave range reported in sleep-associated CSF measurements [11]. The point is not the exact value of a single estimate, but the scale separation it implies: frequencies of order 1 Hz lie well above the poroelastic passband relevant for deep bulk transport.

To test robustness, we varied the main physiological parameters across a plausible range (Table 1). Even when accounting for higher confined moduli for near-incompressible tissue, the resulting cut-off frequencies remain confined to the slow-wave band (∼ 0.02–0.18 Hz).

**Table 1:** Sensitivity analysis of the poroelastic cut-off frequency (*r*_*c*_). Across plausible physiological parameter ranges, the cut-off remains in the slow-wave regime and well below the cardiac band.

| Parameter regime | Permeability<br>($k$) | Confined modulus<br>($M$) | Spacing<br>($L$) | Cut-off<br>($r_c$) |
| --- | --- | --- | --- | --- |
| Nominal baseline | $10^{-14} \text{ m}^2$ | 3 kPa | $300 \mu\text{m}$ | **0.053** Hz |
| Stiffer matrix | $10^{-14} \text{ m}^2$ | 10 kPa | $300 \mu\text{m}$ | **0.177** Hz |
| More compliant /<br>sparser matrix | $2 \times 10^{-14} \text{ m}^2$ | 1 kPa | $400 \mu\text{m}$ | **0.020** Hz |

Under the model assumptions, this establishes a robust mechanical low-pass filter. High-frequency forcing is therefore expected to be strongly attenuated as a driver of deep bulk flow, even if it remains important for local perivascular or interstitial agitation. The corresponding low-pass response and its physiological frequency range are shown in Fig. 2.

**Figure 2:**
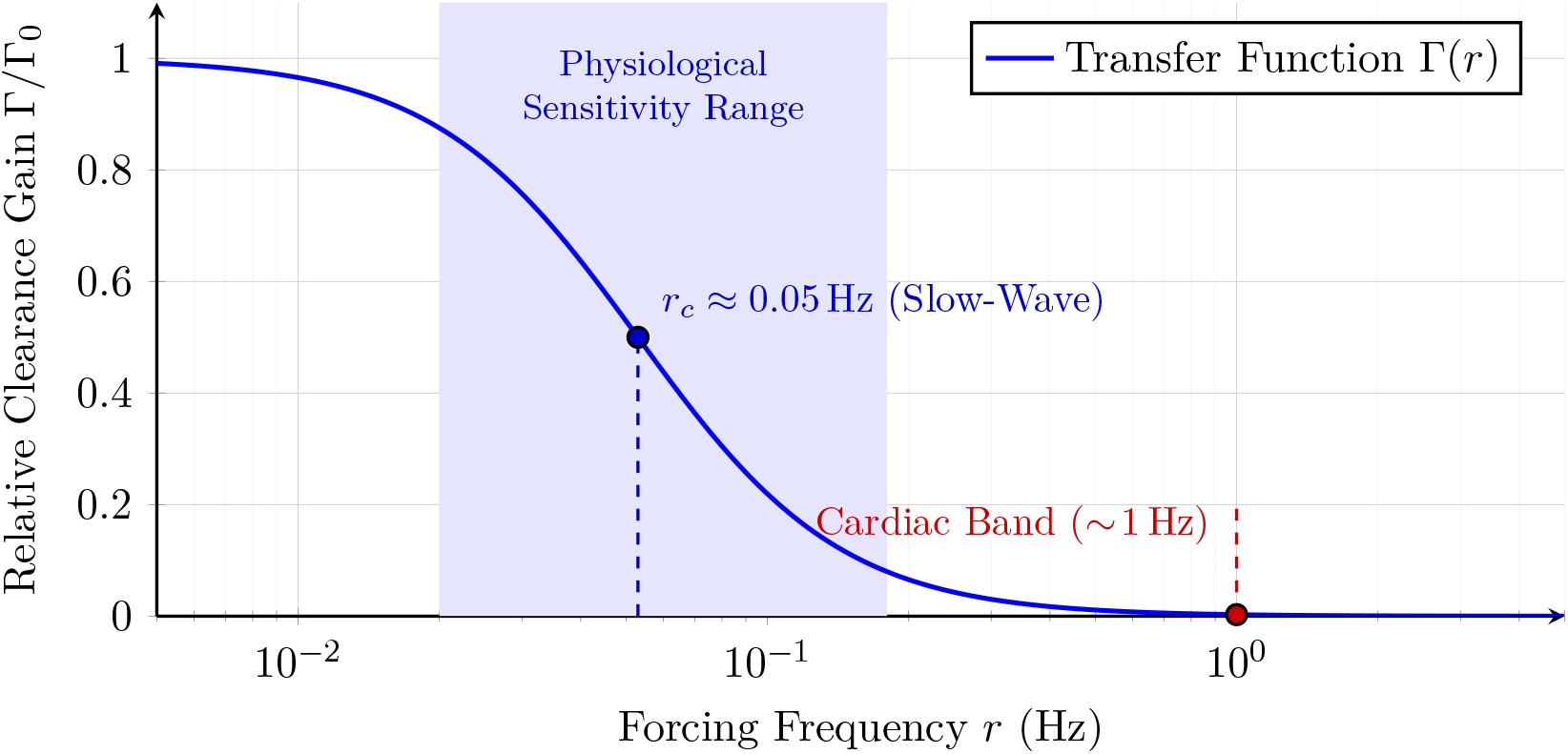
Poroelastic low-pass filtering selects the slow-wave regime for bulk clearance. Lorentzian transfer function Γ(*r*) = Γ_0_[1 + (*r*/*r*_*c*_)^2^]^−1^ shown on a logarithmic frequency axis. The nominal poroelastic cut-off frequency *r*_*c*_ is indicated near 0.05 Hz. Frequencies in the cardiac band (∼ 1 Hz) lie far above the passband and are therefore strongly attenuated as a driver of deep bulk transport.

### 3. Two fluid-dynamical regimes: wakeful boundary forcing and sleep clearance

A purely mechanical account specifies which frequencies and synchrony patterns are favored by tissue transmission, but it does not on its own provide a formal source model for the vascular forcing. To address this, our framework separates the problem into source generation and mechanical transmission (Fig. 3).

**Figure 3:**
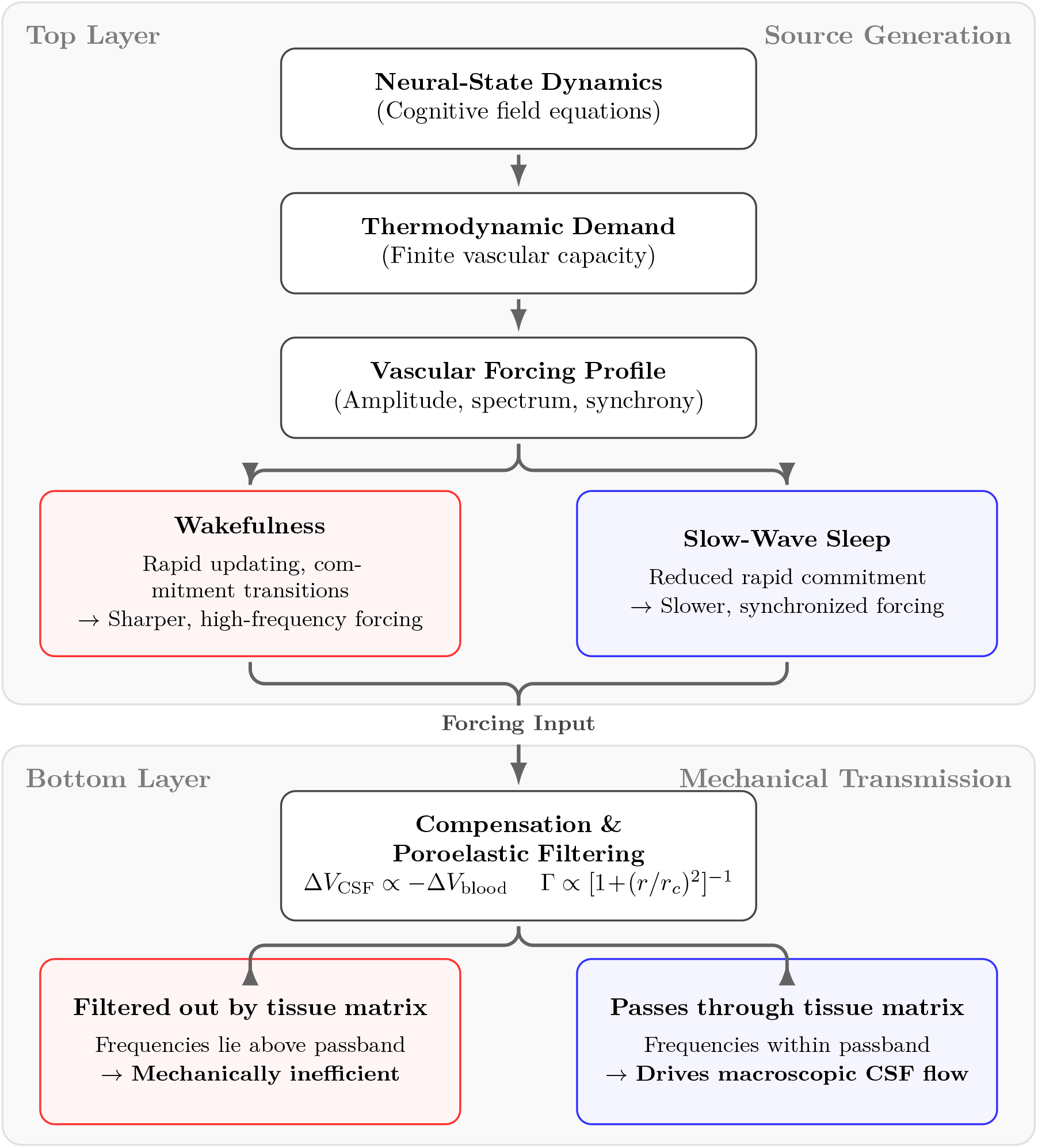
Two-layer account of sleep-associated clearance. The framework separates the problem into source generation and mechanical transmission. **Top layer:** Cognitive field dynamics generate state-dependent vascular forcing through thermodynamic demand. Wakefulness produces sharp, high-frequency transients, while sleep permits slower, synchronized oscillations. **Bottom layer:** This forcing drives intracranial fluid displacement but must pass through the poroelastic tissue matrix. Wakeful forcing is attenuated as it lies above the passband, whereas slow-wave sleep forcing is efficiently transmitted to drive macroscopic CSF flow.

To ground the forcing dynamics in cognitive theory, we represent neural states as probability distributions over competing hypotheses [19]. In this framework, the metabolic cost of state-change is intrinsically coupled to the information geometry of the update [20]. During wakefulness, the system must repeatedly perform fast perceptual, cognitive, and sensorimotor updating. To understand why this matters mechanically, we must introduce a key geometric distinction: the thermodynamic cost of cognitive state change is not evaluated in a single fixed geometry. During wakefulness, the system must repeatedly perform fast perceptual, cognitive, and sensorimotor updating. To understand why this matters mechanically, we must introduce a key geometric distinction: the thermodynamic cost of cognitive state change is not evaluated in a single fixed geometry.

In the companion framework, the effective Bures–Wasserstein state-space geometry changes with uncertainty scale. In the high-uncertainty bulk, distributed inference is governed by Wasserstein transport geometry; near the low-uncertainty boundary, commitment-like compression is governed by Bures covariance geometry. Accordingly, the local demand should be understood schematically as regime dependent,

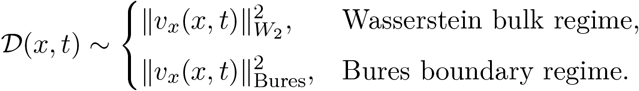

In the present paper, this distinction is used qualitatively to characterize differences in spectral sharpness and synchronization, rather than as an independently validated empirical partition of neural states. This distinction matters mechanically because bulk wandering in the Wasserstein regime is transport-like and comparatively smooth, whereas boundary approach in the Bures regime is compression-like and thermodynamically sharper. When rapid resetting repeatedly drives the system toward the low-uncertainty Bures boundary during wakefulness, the resulting demand profile is not a smooth sinusoidal modulation, but a sequence of relatively sharp, compression-dominated boundary-approach events.

This matters for fluid transport for two reasons. First, the dominant timescale of such wakeful forcing remains too fast relative to the poroelastic cut-off *r*_*c*_, so the tissue strongly attenuates its ability to generate deep bulk flow. Second, because a sharp transient contains more high-frequency harmonic power than a smooth oscillation, the Wasserstein–Bures regime change strengthens the filtering effect: boundary-spiking demand places even more spectral weight above the poroelastic passband than a purely sinusoidal driver at the same base frequency. Wakefulness is therefore mechanically inefficient for bulk clearance not only because it is fast, but because it is dominated by spectrally sharp boundary-approach events.

In this picture, wakefulness need not be described as a collection of small local micro-pumps. That formulation risks suggesting a true clearance mechanism homologous to the slow-wave mode. The more precise claim is that wakefulness generates repeated vascular and fluid displacement attempts, including events that may be spatially widespread or effectively global, but these remain too brief, too sharp, and too high in frequency to survive tissue filtering as coherent bulk clearance.

Slow-wave sleep accesses a different regime. When rapid commitment and continuous sensorimotor correction are suspended, the system can remain for longer intervals in the high-uncertainty Wasserstein bulk. The associated thermodynamic demand is then smoother and can organize into slow, synchronized oscillations across extended regions. Because these oscillations fall within the poroelastic passband, the same Bures–Wasserstein metric-induced volume changes that are mechanically ineffective during wakefulness can now drive macroscopic CSF motion. The extended quiescent interval between successive slow-wave events may also serve a mechanical function, providing the poroelastic matrix sufficient time to relax and refill before the next volumetric stroke. The displacement equation takes the large-scale form

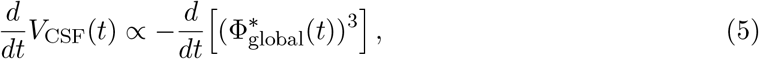

where 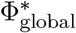 denotes the synchronized effective volume modulation.

The physical interpretation is that sleep does not merely reduce brain activity. It changes the geometry and temporal organization of the forcing. Wakefulness repeatedly drives the system into a metabolically sharp Bures boundary regime whose spectral content is mostly rejected by the tissue. Sleep permits longer residence in the smoother Wasserstein bulk and allows the resulting vascular-volume oscillation to become globally synchronized at a frequency that the tissue can transmit.

### 4. Quantitative predictions: boundary-sensitive clearance and spectral truncation

The framework yields falsifiable predictions that connect geometric state dynamics to measurable fluid and hemodynamic observables.

#### Boundary-sensitive load dependence of sleep clearance

Because displaced CSF volume per cycle scales with the amplitude of the synchronized volumetric factor (Φ^*^)^3^, the model predicts that sleep following a period of high cognitive load should produce larger macroscopic CSF pulsations than sleep following rest, consistent with the observed increase in slow-wave activity and metabolic demand following prolonged wakefulness [26, 27]. The geometric framework sharpens this statement beyond a generic “more effort, more waste” principle. What matters is not only the total amount of processing, but how strongly wakefulness repeatedly drives the system toward the low-uncertainty Bures boundary.

Tasks dominated by shallow exploration in the Wasserstein bulk should generate comparatively smooth and moderate demand. By contrast, tasks requiring repeated high-precision commitment, rapid decision closure, or strong error suppression should generate sharper, more compression-dominated demand transients because they repeatedly enter the Bures boundary regime.

Physiologically, this follows because prolonged wakeful boundary forcing is expected to leave the substrate closer to its effective capacity limit at sleep onset. Elevated waking demand drives greater accumulation of metabolic waste, which physically lowers the effective autoregulatory capacity *Q*_max_ at sleep onset. In the constrained regime,

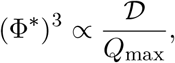

so a depressed *Q*_max_ or elevated residual demand implies a larger slow-wave volumetric oscillation and hence greater CSF displacement per cycle.

#### BOLD-first / CSF-second ordering

The model predicts that the temporally leading event in a clearance episode should be a synchronized vascular-volume increase, with the CSF pulse following as a delayed mechanical consequence. In other words, the large slow-wave CSF inflow should be preceded by a hemodynamic event, not accompanied by an independent fluid driver.

This follows directly from the causal chain

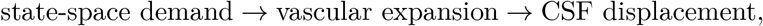

together with the poroelastic filtering of the tissue response.

#### Two-compartment transport discrepancy and tracer dispersion

The poroelastic filter natively predicts a striking mechanical discrepancy between the transport of deep endogenous waste and the dispersion of superficial exogenous tracers. This follows directly from the role of hydraulic permeability (*k*) in the cut-off frequency, *r*_*c*_ ∝ *k*. In the dense interstitial matrix where cognitive byproducts are generated, permeability is strictly limited (*k* ≈ 10^−14^m^2^), yielding the nominal slow-wave passband *r*_*c*_ ≈ 0.05 Hz. At waking cardiac frequencies (*r* ≈ 1 Hz), the transmitted fluid response Γ is strongly suppressed:

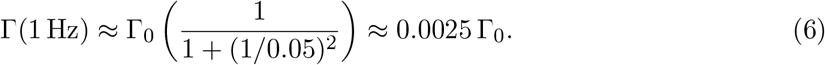

The high-frequency waking drive is attenuated by over 99%, trapping deep interstitial waste. Conversely, exogenous tracers are typically injected into perivascular spaces. These fluid channels possess orders-of-magnitude higher permeability, shifting their local cut-off frequency well above the cardiac band (*r*_*c*_ ≫ 1 Hz). For these superficial compartments, the high-frequency waking drive is fully transmitted:

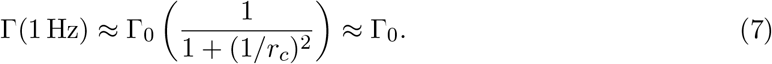

Consequently, the model predicts that the waking state will simultaneously fail to clear deep interstitial waste while generating sufficient local perivascular agitation to rapidly scatter exogenous tracers.

#### Spectral truncation scaling law for resting-state networks

The framework also provides a geometric and quantitative mechanism for the spectral alterations observed in resting-state fMRI during physiological aging and neurodegeneration [28, 29]. During quiet wakefulness, the system is assumed to spend most of its time wandering stochastically in the high-uncertainty, bulk-like regime, consistent with dynamical-systems models of resting-state activity as a continuous stochastic exploration of state space [30].

For two nodes separated by a baseline topological distance *d*, the characteristic stochastic traversal time, or coherence time 𝒯_coh_, scales quadratically with effective geometric distance:

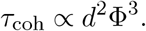

Substituting the constitutive scaling law for metric expansion in the capacity-limited regime gives

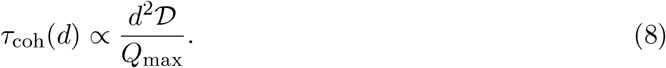

Consequently, the intrinsic frequency *f*_*d*_ of a stochastically coupled subnetwork scales as the inverse of this coherence time:

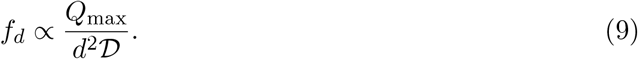

This equation implies two linked consequences. First, local subnetworks with smaller effective separation *d* naturally support higher frequencies. Second, there is a metabolic truncation scale for large networks. Requiring a minimum viable coherence frequency *f*_min_ gives a maximum sustainable network diameter

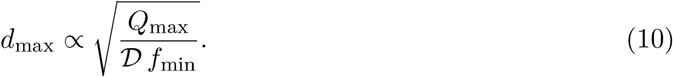

As vascular capacity *Q*_max_ declines, large-scale low-frequency modes become unsustainable and are lost first. Networks larger than *d*_max_ are therefore predicted to fragment metabolically before smaller local networks do.

The Wasserstein–Bures distinction also refines this prediction. Quiet resting-state fluctuations should be governed primarily by Wasserstein bulk wandering, validating the divergence-like scaling above. However, occasional boundary-approach excursions during wakefulness remain possible. Because those excursions are disproportionately costly in the Bures regime, they should generate a heavy-tailed demand distribution superimposed on the bulk fluctuations.

## Discussion

The question of why sleep is mechanically privileged for large-scale CSF dynamics remains unresolved because each proposed driver captures part of the physiology but not the full state dependence of the phenomenon [4].

Arterial pulsation models [5] provide a continuously available forcing source, but under the coarse-grained poroelastic estimates developed here—and consistent with fluid-dynamical models of perivascular transport [7]—much of that forcing lies above the effective passband for deep bulk transport. Respiratory models [8] occupy a more favorable frequency range but remain present during both wake and sleep, making them insufficient on their own to explain the strong sleep dependence of large-scale CSF dynamics. Vasomotion [9, 10] approaches the relevant band more closely, but is often spatially local and heterogeneous rather than globally phase-locked. On this view, the key issue is not whether these candidate drivers exist, but which classes of forcing are mechanically compatible with large-scale transmission through brain tissue.

The present framework offers a way to organize these observations. It does not deny that arterial, respiratory, and vasomotor oscillations contribute to intracranial fluid dynamics. Instead, it proposes a selection principle: forcing modes are expected to contribute most effectively to large-scale bulk transport when they are both sufficiently slow and sufficiently synchronized to survive tissue filtering. Slow-wave sleep is special in this view not because it introduces an entirely new fluid driver, but because it permits brain-wide coordination in a frequency range that the tissue can actually transmit.

This selection principle directly addresses the mechanical foundations of the glymphatic system hypothesis [1, 14]. While the existence of perivascular pathways and the critical role of astroglial aquaporin-4 (AQP4) channels in facilitating fluid exchange are widely supported, the dominant driving force of this network remains contested. The original glymphatic model posits arterial pulsation as an important mechanical driver. However, our poroelastic filter analysis suggests that while high-frequency pulsations may support local perivascular exchange, they are likely to be mechanically inefficient as drivers of deep interstitial bulk transport. In that context, the present framework proposes a complementary macroscopic source of forcing—slow, thermodynamically constrained vascular volume modulation—operating in a frequency range more compatible with tissue-level transmission. On this view, the geometric state-dynamics developed here offer a candidate mechanical basis for the slow, high-amplitude forcing that sleep-related glymphatic transport may require.

Recent computational models have independently shown that slow vascular volume changes are mechanically more effective than high-frequency pulsations at driving convective fluid motion through a poroelastic brain matrix [31]. What the present framework adds to these mechanical accounts is a candidate upstream source model for that relevant forcing. In our formulation, synchronized blood-volume modulation is interpreted as the physiological expression of substrate-constrained neural-state dynamics. The central claim is therefore not that sleep introduces a new driver of intracranial fluid motion, but that slow-wave sleep places the existing neurovascular machinery into a forcing regime that is mechanically compatible with large-scale transmission through brain tissue.

Beyond the biomechanics, the framework suggests that the state dependence of sleep-associated CSF dynamics may reflect a deeper link between neural-state change and substrate thermodynamics. In the companion formalism [18, 32], wakeful commitment is modeled as repeated approach to a low-uncertainty boundary regime, whereas slow-wave sleep permits longer residence in a smoother, bulk-like regime. In the present paper, that distinction is given a mechanical reading: wake-like dynamics generate sharp, high-frequency vascular forcing that is poorly matched to the poroelastic passband, whereas slow-wave sleep permits slower and more synchronized forcing that can be more effectively transmitted. On this view, the sleep–wake contrast is not merely a difference in overall activity level, but a difference in the spectral and geometric organization of the forcing delivered to the tissue.

This geometric organization can also be interpreted more formally within the companion framework [18, 32]. In that reading, the conformal response Φ^*α*^ expresses a balance between thermodynamic demand and effective substrate capacity. During wakefulness, the resulting forcing is modeled as sharper and more spectrally concentrated, making it poorly matched to the poroelastic passband. During slow-wave sleep, that mismatch is reduced: the forcing becomes smoother and more synchronized, allowing more effective large-scale transmission through the tissue.

The same framework also offers one possible way to reconcile a central experimental tension in the literature: wakefulness may be inefficient for deep interstitial bulk transport while still permitting rapid dispersion of superficial exogenous tracers. In the present model, this difference follows from compartment-specific permeability. High-frequency forcing can be strongly attenuated in the dense interstitial matrix while remaining mechanically effective in more weakly resistive perivascular spaces. On this view, the apparent disagreement between sleep-dependent endogenous clearance and wake-associated tracer dispersion need not reflect incompatible physics, but different transport regimes of the same poroelastic system.

Several caveats are important. First, the link between cognitive demand and the squared state-space velocity, 𝒟 ∝ ∥*v*_*x*_∥^2^, is used here as a constitutive physiological assumption rather than a formal theorem, although it is motivated by the kinetic-energy formulation of optimal transport [20, 18]. Second, the interpretation of (Φ^*^)^3^ as a direct proxy for local cerebral blood-volume change is an idealization. Third, the poroelastic calculation is deliberately coarse-grained to identify the controlling frequency scale and passband logic, not to replace detailed, spatially resolved biomechanical modeling of tissue anisotropy and macroscopic fluid dynamics [33]. Fourth, the wake–sleep geometric contrast relies on the formal regime change derived in the companion framework [18, 32]; evaluating this foundation requires joint consideration of those mathematical theories. Most importantly, the present model addresses the transmission of oscillatory fluid forcing and its compatibility with deep bulk transport; it does not by itself constitute a full quantitative theory of net solute clearance.

Even with those limitations, the framework has a useful unifying power. It links cognitive state dynamics, vascular expansion, intracranial volume compensation, tissue filtering, and sleep-associated clearance in a single mechanical narrative. In this narrative, slow-wave sleep is not merely a state in which clearance-related CSF dynamics happen to occur. Rather, the suspension of rapid distributed commitment cycles permits the brain to enter a slow synchronized regime that is mechanically compatible with whole-brain fluid transport.

## Methods

### Poroelastic filtering model and sensitivity analysis

We model the tissue as a poroelastic medium in the sense of Biot theory [21, 22, 23]. The physical properties of this interstitial matrix, including its high tortuosity and strictly limited hydraulic permeability, are well established in the experimental and biomechanical literature [24]. The cut-off scale *r*_*c*_ follows from the poroelastic consolidation time:

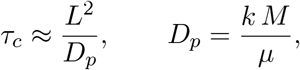

where *L* is the relevant drainage length scale, *k* is hydraulic permeability, *M* is the confined (oedometric) modulus, and *µ* is fluid viscosity. For nearly incompressible biological tissue, *M* is typically larger than Young’s modulus (*E*). We utilize a conservative lower bound of *M* ≈ *E* ≈ 3 × 10^3^ Pa. The associated cut-off is

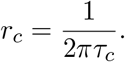

Using conservative literature values for cortical tissue,

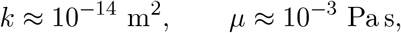

we obtain *D*_*p*_ ≈ 3 × 10^−8^ m^2^*/*s. Taking the relevant drainage length to be the approximate spacing between penetrating arterioles, *L* ≈ 300 *µ*m [34, 35], gives

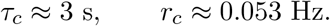

To ensure the filter is robust to parameter variations, we calculated *r*_*c*_ across a physiological sensitivity range (Table 1).

### Cognitive field source model

The mechanical analysis in the main text specifies which classes of vascular forcing are expected to be compatible with deep bulk transport. To model how such forcing arises from neural-state dynamics, we introduce a constrained state-space description conceptually allied with free-energy formulations of brain function [19] and the extended state-space model [32].

Let *x*(*t*) denote the effective neural state and *v*_*x*_(*x, t*) the associated state-space velocity field. The instantaneous thermodynamic demand associated with state updating is modeled as 𝒟 (*x, t*) ∝ ∥*v*_*x*_(*x, t*)∥^2^. The substrate is assumed to have finite local perfusion capacity *Q*_max_(*x, t*). In the constrained regime, increasing demand relative to available capacity induces a conformal modification of the effective Bures–Wasserstein geometry [18] through a conformal response factor Φ^*^(*x, t*):

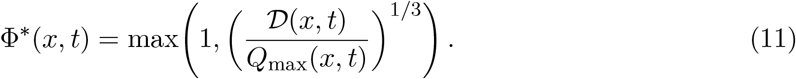

The term (Φ^*^)^3^ is treated as a constitutive proxy for local cerebral blood-volume modulation: *CBV* (*x, t*) ∝ (Φ^*^(*x, t*))^3^.

### State-regime interpretation

To compare wakefulness and slow-wave sleep, we use the geometric regime distinction developed in companion work [18, 32]. In high-uncertainty regions of state space, updating is described by Wasserstein transport geometry; near low-uncertainty, commitment-like states, the dynamics are boundary-weighted and described by Bures covariance geometry. Wake-like forcing is accordingly modeled as a regime enriched in rapid resetting and commitment-like transitions (*z* → *z*_0_), generating sharper vascular transients. Slow-wave-like forcing is modeled as a regime in which such transitions are reduced, allowing smoother oscillations to emerge.

## Supplementary Information

### S1 Generalization to non-Euclidean vascular topologies

In the main text, the volumetric Bures–Wasserstein metric response is derived using an isotropic Euclidean scaling exponent *α* = 3, leading to the effective Bures–Wasserstein metric

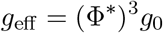

and the scaling law

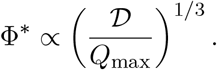

This assumption models the microvascular bed as a space-filling local expansion.

Real cerebrovascular networks, however, are highly branched and are more accurately described by non-Euclidean or fractal morphologies [36]. In such a case, the geometric response need not occupy a full three-dimensional Euclidean expansion volume. If the effective dimensionality of the vascular response is *D*_*f*_ *<* 3, the constitutive law generalizes to

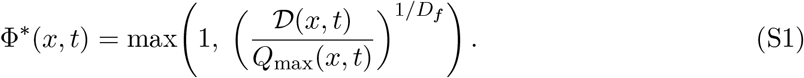

Because *D*_*f*_ *<* 3, the exponent 1*/D*_*f*_ exceeds the Euclidean exponent 1*/*3. The consequence is an accelerated metric stiffening effect: for a given capacity deficit D*/Q*_max_ *>* 1, a fractal substrate undergoes steeper conformal stretching than a homogeneous Euclidean volume. The Euclidean case used in the main text should therefore be read as a conservative lower-bound model for the magnitude of the response, not as the only admissible geometry.

### S2 Projection-induced memory and hemodynamic delay

Standard hemodynamic models, such as the classic Balloon model, often attribute the characteristic delay of the BOLD response primarily to vascular compliance [37, 38]. In the broader geometric framework motivating the present paper, there is potentially a second source of effective memory: projection over hidden state variables.

The full system is assumed to be Markovian only on an extended state space containing both the visible hypothesis variables and one or more hidden uncertainty or substrate coordinates. Let the instantaneous geometric demand be

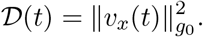

If the conformal factor relaxes with a finite timescale 𝒯according to

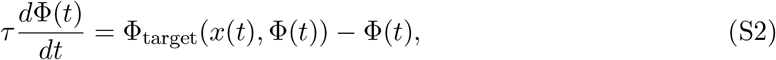

then observing only a projected macroscopic variable induces an effective memory kernel, even when the underlying extended dynamics are Markovian.

To leading order, this projection yields a coarse-grained effective demand of the form

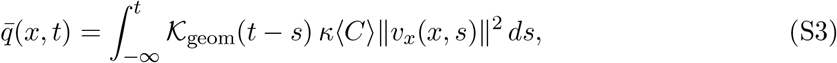

where 𝒦_geom_ is an exponential-like relaxation kernel capturing the hidden geometric dynamics. If the resetting operator acts at a rate *r* (e.g., the ∼ 1 Hz cardiac pulse), the effective decay time scales as 𝒯_eff_ ≈ (*λ* + *r*)^−1^, where *λ* governs the local drift of the uncertainty coordinate. The observed hemodynamic response may then be represented schematically as

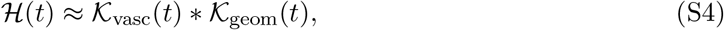

where 𝒦_vasc_ is the purely vascular component and 𝒦_geom_ is the projection-induced geometric component.

The present CSF paper does not require this decomposition for its main argument. We record it here because it suggests a further empirical prediction: tasks differing mainly in uncertainty-structure rather than total energetic demand may still exhibit different fitted hemodynamic latencies. This establishes a formal double dissociation: pharmacological or physiological manipulations targeting vascular compliance should primarily alter K_vasc_, whereas cognitive manipulations of uncertainty and task-resetting rates should primarily modulate the projection kernel 𝒦_geom_.

### S3 Field-theoretic origin of the macroscopic source-to-transport chain

The purpose of this section is to show how the macroscopic volume-compensation relation can be embedded within a broader formal framework linking information-geometric state dynamics to continuum mechanics. While the biomechanical results in the main text are derived phenomenologically, this field-theoretic approach—conceptually allied with the study of substrate-constrained mathematical triads [18]—provides a candidate first-principles derivation of the vascular source term.

In this formulation, the neurovascular substrate satisfies a self-consistency condition at the cognitive condensation boundary. The conformal response Φ^α^ is interpreted as a geometric back-reaction to the thermodynamic demand 𝒟. This establishes a cognitive-vascular “equation of state” where the realized vascular strain must balance the metabolic dissipation rate dictated by the gradient flow on state space. Within this geometric-thermodynamic duality, the Bures–Wasserstein regime change acts as a mechanical engine: the transition from distributed deliberation (Wasserstein bulk) to sharp commitment (Bures boundary) dictates the spectral forcing profile delivered to the poroelastic tissue matrix.

#### Conformal response as an effective active strain field

In the companion field-theoretic framework, cognitive demand induces a local Bures–Wasserstein metric response on the neural manifold M. The effective metric is taken to be a conformal deformation of the resting background metric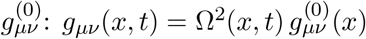, where Ω^2^(*x, t*) = (Φ^*^(*x, t*))^3^. To connect this geometric response to continuum mechanics, we introduce a small-deformation active strain-like tensor: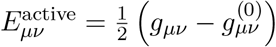. Its trace is proportional to the conformal volume modulation ((Φ^*^)^3^ − 1).

#### Coupling to poroelastic mass conservation

We next embed this active-strain source in a Biot-type poroelastic description of fluid transport. Let *w*^*µ*^ denote the covariant fluid flux vector and *p* the pore pressure. Conservation of fluid mass requires: 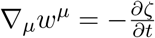 where 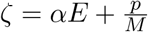. Substituting the active geometric strain trace gives:

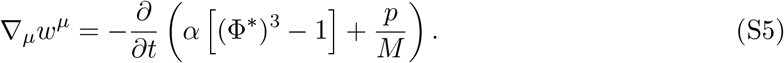

Under this constitutive identification, the temporal evolution of the conformal factor acts as the scalar source term driving the fluid flux field *w*^*µ*^. The transfer function Γ(*r*) in the main text can be interpreted as the effective Green’s-function response of this coupled poroelastic system.

#### Global volume compensation as a boundary-flux constraint

The Monro–Kellie doctrine emerges as an effective boundary-flux constraint on the intracranial domain V. Integrating the mass balance and applying the divergence theorem yields:

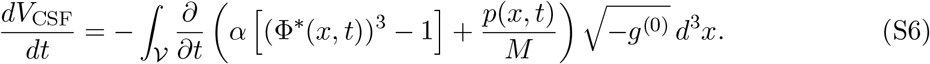

The macroscopic Monro–Kellie pump is thus the global conservation consequence of a thermodynamically driven geometric source field evolving in a poroelastic medium under constrained boundary flux.

## Notes

### Competing Interest Statement

The authors have declared no competing interest.

